# Rictor/TORC2 mediates gut-to-brain signaling in the regulation of phenotypic plasticity in *C. elegans*

**DOI:** 10.1101/189613

**Authors:** Michael P. O’Donnell, Pin-Hao Chao, Jan E. Kammenga, Piali Sengupta

## Abstract

Animals integrate external cues with information about internal conditions such as metabolic state to execute the appropriate behavioral and developmental decisions. Information about food quality and quantity is assessed by the intestine and transmitted to modulate neuronal functions via mechanisms that are not fully understood. The conserved Target of Rapamycin complex 2 (TORC2) controls multiple processes in response to cellular stressors and growth factors. Here we show that TORC2 coordinates larval development and adult behaviors in response to environmental cues and feeding state in the bacterivorous nematode *C. elegans*. During development, pheromone, bacterial food, and temperature regulate expression of the *daf-7* TGF-β and *daf-28* insulin-like peptide in sensory neurons to promote a binary decision between reproductive growth and entry into the alternate dauer larval stage. We find that TORC2 acts in the intestine to regulate neuronal expression of both *daf-7* and *daf-28,* which together reflect bacterial-diet dependent feeding status, thus providing a mechanism for integration of food signals with external cues in the regulation of neuroendocrine gene expression. In the adult, TORC2 similarly acts in the intestine to modulate food-regulated foraging behaviors via the PDFR-1 neuropeptide receptor. We also demonstrate that genetic variation affects food-dependent larval and adult phenotypes, and identify quantitative trait loci (QTL) associated with these traits.Together, these results suggest that TORC2 acts as a hub for communication of feeding state information from the gut to the brain, thereby contributing to modulation of neuronal function by internal state.

**AUTHOR SUMMARY:** Decision-making in all animals, including humans, involves weighing available information about the external environment as well as the animals’ internal conditions. Information about the environment is obtained via the sensory nervous system, whereas internal state can be assessed via cues such as levels of hormones or nutrients. How multiple external and internal inputs are processed in the nervous system to drive behavior or development is not fully understood. In this study, we examine how the nematode *C. elegans* integrates dietary information received by the gut with environmental signals to alter nervous system function. We have found that a signaling complex, called TORC2, acts in the gut to relay nutrition signals to alter hormonal signaling by the nervous system in *C. elegans*. Altered neuronal signaling in turn affects a food-dependent binary developmental decision in larvae, as well as food-dependent foraging behaviors in adults. Our results provide a mechanism by which animals prioritize specific signals such as feeding status to appropriately alter their development and/or behavior.

## INTRODUCTION

Animals integrate complex and varying environmental stimuli in the context of their past experience and current conditions to execute the appropriate developmental and behavioral response. A particularly critical environmental variable is the availability of food. Nutritional experience regulates developmental trajectories and behavioral decisions, and modulates physiological state across phyla (1-5). In the majority of animal taxa, the primary site of nutrient uptake is through the digestive system. The intestine not only plays a role in nutrient acquisition, but also transmits signals conveying nutrient status to multiple tissues including the nervous system (6-10). The importance of gut-to-brain signaling in the regulation of both developmental and behavioral traits has now been demonstrated in several systems, and a subset of the underlying biochemical pathways has been identified (11-17). However, how nutrient status is assessed and integrated with other cues to modulate development and behavior remains to be fully described.

The bacterivorous nematode *Caenorhabditis elegans* displays a complex repertoire of sensory-driven developmental and behavioral traits, many of which are regulated by the animal’s feeding status (18-22). During early larval development, cues such as high temperature and high pheromone levels (reflecting population density) promote entry of juvenile *C. elegans* into the alternate dauer developmental stage (23-27). The salience of dauer-promoting cues is modulated by food availability (20, 28, 29). Abundant food suppresses the dauer-promoting effects of temperature and pheromone, and conversely, food restriction enhances their effects (20, 28). Thus, analysis of dauer formation provides an opportunity to explore the mechanisms by which nutrient signals are integrated with other sensory stimuli to regulate a binary developmental decision.

Sensory cues target two parallel neuroendocrine pathways to regulate dauer formation (30). Pheromone downregulates expression of the DAF-7 TGF-β ligand, and alters the expression of multiple insulin-like peptides (ILPs) including the DAF-28 ILP in sensory neurons (28, 31-35). Expression of both *daf-7* and *daf-28* is also strongly regulated by feeding status (28, 31, 36-38), suggesting that multiple stimuli converge on the modulation of expression of these neuroendocrine genes to regulate dauer entry.Whether food availability is encoded by altered external sensory inputs and/or internal nutrient status to regulate neuroendocrine gene expression is not fully understood.

In addition to regulating developmental plasticity, perception of food as well as assessment of internal metabolic state also modulates behaviors and physiology of *C. elegans* adults [eg. (3, 11, 18, 19, 39)]. On a bacterial lawn, well-fed wild-type adult animals spend the majority of their time in a low activity dwelling state whereas adults that are unable to efficiently absorb nutrients exhibit higher-activity roaming, or exploratory behavior (21, 40, 41). The transition between these locomotory states appears to be largely regulated by two antagonistic neuromodulatory pathways involving the PDF neuropeptide and serotonin, although additional molecules including DAF-7 TGF-β have also been implicated in this process (21, 42, 43). Internal metabolic state has been suggested to regulate this behavioral transition (21), suggesting that gut-to-brain signaling plays an important role in regulating adult exploratory behavior.

The mechanistic Target of Rapamycin (mTOR) pathway consists of the highly conserved TORC1 and TORC2 molecular complexes that act as cell-intrinsic sensors and regulators of metabolic status in multiple species (44-46). The well-studied TORC1 complex responds to nutrient and growth factor signals and regulates a diverse array of cell-biological processes associated with growth and proliferation (47-49). Although the TORC2 complex also responds to growth factors and regulates a partially overlapping set of cellular processes with TORC1, the regulation and targets of this complex are less well-characterized (50-52). In *C. elegans,* TORC2 activity regulates physiological processes such as growth rate, lipid production, and longevity and has been suggested to act as a sensor for internal energy status (Figure 1A) (11, 53-55). Interestingly, the effects of TORC2 on lifespan appear to be modulated by bacterial diet and growth temperature (11, 54, 56), and TORC2 signaling was also shown to regulate diet-dependent effects on behavioral traits (55). TORC2 appears to act either in the intestine or in neurons via activation of the transcription factors DAF-16/FoxO or SKN-1/Nrf (54, 55) to regulate different processes (Figure 1A).

**Figure 1.**
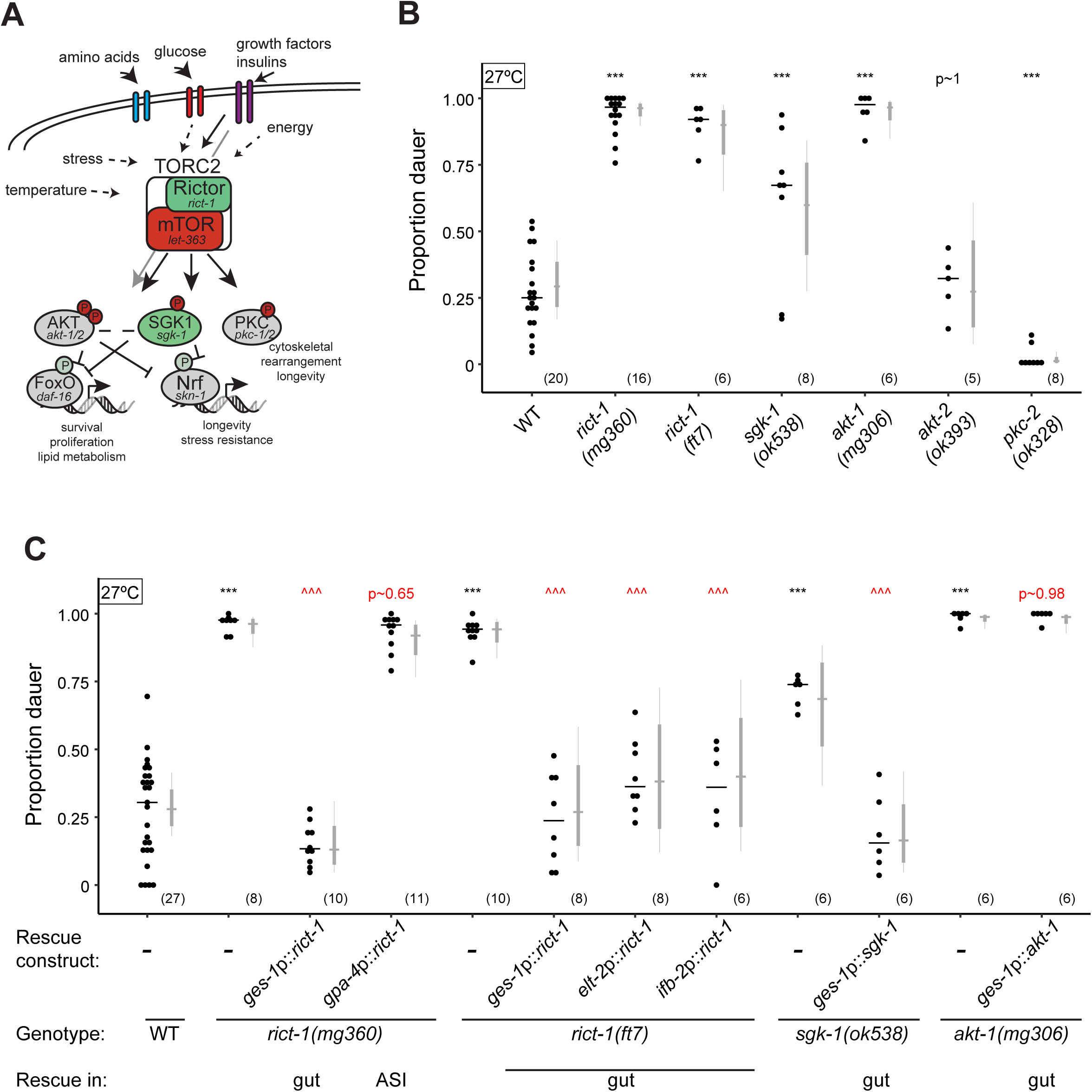
The TORC2 signaling pathway suppresses high temperature-induced dauer formation. **A)** The TORC2 signaling pathway in *C. elegans* (11, 53-55). Genes encoding conserved components of the pathway are indicated. **B-C)** Dauers formed by animals of the indicated genotypes at 27°C. Each black dot indicates the proportion of dauers formed in a single assay. Horizontal black bar indicates median. Light gray thin and thick vertical bars at right indicate Bayesian 95% and 75% credible intervals, respectively. Numbers in parentheses below indicate the number of independent experiments with at least 25 and 9 animals each scored for non-transgenic and transgenic animals, respectively. For transgenic rescues, data are combined from 2 independent lines, with the exception of *ges-1p* constructs, in which one line was analyzed. *** - *P*<0.001 compared to wild-type (ANOVA with Dunnett-type multivariate-t adjustment for 1B and Tukey-type multivariate-t adjustment for 1C); ^^^ - *P*<0.001 compared to corresponding mutant animals. *P*-values of differences in means relative to wild-type (B) and corresponding mutant animals (C) are indicated in black and red, respectively.

Here we report that intestinal TORC2 activity is required for food-dependent modulation of dauer entry and exploratory behavior in *C. elegans.* We show that the essential TORC2 component *rict-*1/Rictor acts in the intestine to promote neuronal expression of the *daf-7* TGF-β and the *daf-28* ILP genes. We find that food-dependent TORC2 signals acting in parallel to temperature and pheromone signaling converge, and are integrated, at the level of regulation of *daf-7* and *daf-28* expression in sensory neurons to modulate dauer formation. Consistent with decreased detection or utilization of food signals in the absence of Rictor function, *rict-1* mutants exhibit enhanced dauer formation in response to both temperature and pheromone. We also implicate intestinal TORC2 signaling in the regulation of food-dependent adult foraging decisions. Via analyses of dauer formation and adult foraging behavior in *C. elegans* wild isolates, we show that these traits are genetically separable, and identify quantitative trait loci (QTL) that contribute to trait variation between two strains. Our results indicate that intestinal TORC2 plays a critical role in conveying internal nutrient state information to the nervous system, thereby allowing animals to modulate both developmental and behavioral responses to environmental cues as a function of their metabolic state.

## RESULTS

### Rictor acts in the intestine to regulate high temperature-induced dauer formation

Given the role of TORC2 in regulating *C. elegans* longevity in a temperature and food-dependent manner (11, 54-56), we asked whether TORC2 also regulates dauer formation – a developmental polyphenism that is also regulated by temperature and food (20, 25, 27). Loss-of-function mutations in *rict-1* Rictor, *sgk-1* SGK1, and *akt-1* AKT/PKB, components of the TORC2 signaling pathway (Figure 1A), did not affect dauer formation in the presence of plentiful OP50 bacteria and without added exogenous pheromone at 25°C [0 dauers for N2 wild-type, *rict-1(ft7), rict-1(mg360), sgk-1(ok538)* and *akt-1(mg306)*; n > 6 assays with >50 animals each). However, mutations in all three molecules, but not in the *akt-2* AKT/PKB or *pkc-2* PKC genes, resulted in significantly increased dauer formation upon a temperature shift to 27°C [high temperature-induced dauer formation or Hid phenotype (27); Figure 1B], suggesting that TORC2 suppresses high temperature-induced dauer formation.

To identify the site of TORC2 activity in regulating dauer formation at high temperatures, we performed tissue-specific rescue experiments. Expression of wild-type *rict-1* cDNA sequences in the intestine using the *ges-1* (57), *elt-2* (58) or *ifb-2* (59) promoters, but not in the ASI sensory neurons using a *gpa-4* promoter (60), robustly rescued the Hid phenotype of both *rict-1(mg360)* hypomorphic and *rict-1(ft7)* putative null mutants (Figure 1C). Similarly, intestinal expression of *sgk-1* rescued the Hid phenotype of *sgk-1(ok538)* mutants (Figure 1C). However, expression of *akt-1* in the intestine failed to rescue the Hid phenotype of *akt-1(mg306)* animals (Figure 1C), suggesting that AKT-1 may act in tissues other than, or in addition to, the intestine to regulate dauer formation at high temperatures. Intestinal TORC2 has been shown to act via SGK-1 and SKN-1 to regulate longevity in a temperature-dependent manner (54). However, we were unable to assess the Hid phenotype of *skn-1* mutants due to their early larval arrest upon growth at 27°C (data not shown). We conclude that TORC2 acts in the intestine, possibly via SGK-1, to regulate high temperature-driven dauer formation.

### Intestinal Rictor regulates neuronal TGF-β and insulin expression

To test whether intestinal TORC2 signaling targets the dauer-regulatory *daf-7* TGF-β and *daf-28* ILP neuroendocrine signaling pathways, we examined the expression of *daf-7* and *daf-28* reporter transgenes in *rict-1* mutants. We found that expression of a stably integrated *daf-7*p::*gfp* reporter transgene (61) was 1.3-2-fold decreased in the ASI neurons of *rict-1* mutant L1 larvae at 27°C (Figure 2A). Consistent with this observation, *daf-7* transcript levels were also reduced in ASI in *rict-1* mutants as assessed via fluorescent *in situ* hybridization (Figure 2B). The reduced expression of *daf-7*p::*gfp* as well as *daf-7* transcript levels in ASI was rescued upon expression of wild-type *rict-1* sequences in the intestine (Figure 2A,B). Overexpression of *daf-7* in ASI using the *srg-47* promoter partly suppressed the Hid phenotype of *rict-1* mutants (Figure 2C). These results suggest that Rictor acts in the intestine to regulate *daf-7* TGF-β mRNA levels in the ASI neurons, and that reduced TGF-β expression in *rict-1* mutants partly underlies their Hid phenotype.

**Figure 2.**
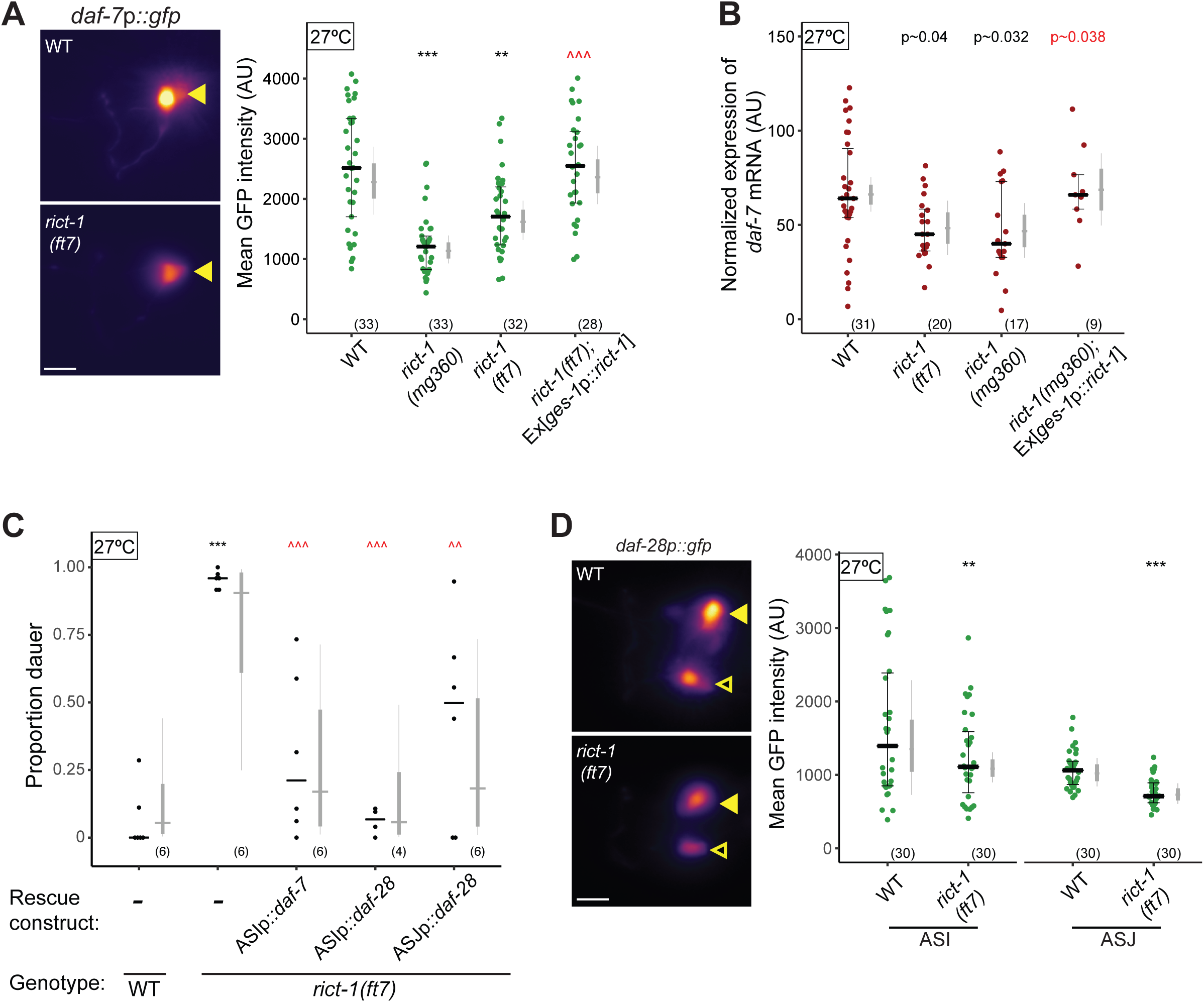
Rictor acts in the intestine to regulate *daf-7* TGF-β and *daf-28* ILP gene expression in sensory neurons. **A,D)** (Left) Representative images of *daf-7*p::*gfp* (A) and *daf-28*p::*gfp* (D) expression in one of the bilateral ASI (A), and ASI and ASJ (D), neurons in animals of the indicated genotypes. Yellow filled and open arrowheads indicate ASI and ASJ soma, respectively.Anterior is at left. Scale bar: 5 μm. (Right) Quantification of *daf-7*p::*gfp* expression in ASI (A) and *daf-28*p::*gfp* expression in ASI and ASJ (D). Each dot is the meanfluorescence intensity in a single animal (2 neurons per animal); numbers in parentheses below indicate the number of animals examined in 3 independent experiments.Horizontal thick bar indicates median. Error bars are quartiles. Light gray thin and thick vertical bars at right indicate Bayesian 95% and 75% credible intervals, respectively. ** and *** - different from wild-type at *P*<0.01 and *P*<0.001, respectively; ^^^ - different from *rict-1(ft7)* at *P*<0.001 (ANOVA with Tukey-type multivariate-t post-hoc adjustment). **B)** Quantification of *daf-7* mRNA levels in ASI assessed via fluorescent *in situ* hybridization. Expression was normalized by subtracting mean background pixel values for each image. Each dot is the fluorescence intensity in a single ASI neuron; numbers in parentheses below indicate the number of neurons examined in 3 independent pooled experiments. Horizontal thick bar indicates median. Error bars are quartiles. Light gray thin and thick vertical bars at right indicate Bayesian 95% and 75% credible intervals, respectively. *P*-values of differences relative to wild-type and *rict-1(mg360)* animals are indicated in black and red, respectively. **C)** Dauers formed by animals of the indicated genotypes at 27°C. Each black dot indicates the average number of dauers formed in a single assay. Horizontal black bar indicates median. Light gray thin and thick vertical bars at right indicate Bayesian 95% and 75% credible intervals, respectively. Numbers in parentheses below indicate the number of independent experiments with at least 36 and 9 animals each scored for non-transgenic and transgenic animals, respectively. Promoters driving expression in ASI andASJ were *srg-47*p and *trx-1*p, respectively. One transgenic line was tested for each condition. *** - different from wild-type at *P*<0.001; ^^ and ^^^ - different from *rict-1(ft7)* at *P*<0.01 and *P*<0.001, respectively (ANOVA with Tukey-type multivariate-t post-hoc adjustment). *P*-values of differences in means relative to wild-type (B) and corresponding mutant animals (C) are indicated in black and red, respectively.

TGF-β and ILP signaling act in parallel to regulate dauer formation (30). Since overexpression of *daf-7* only partially suppressed the *rict-1* Hid phenotype, we considered the possibility that RICT-1 also regulates ILP signaling. Indeed, we observed reduced expression of a *daf-28*p::*gfp* reporter gene in both the ASI and ASJ neurons in *rict-1* mutants (Figure 2D). Similar to our results upon overexpression of *daf-7*, overexpression of *daf-28* in either ASI or ASJ reduced the Hid phenotype of *rict-1* mutants (Figure 2C). Moreover, a mutation in the *daf-16* FOXO transcription factor gene– a key target inhibited by ILP signaling – partly suppressed the Hid phenotype of *rict-1* mutants (Figure S1A). Together, these results indicate that RICT-1 regulates *daf-7* TGF-β and *daf-28* ILP expression in the ASI and ASI/ASJ sensory neurons, respectively, to regulate dauer formation at 27°C.

### Heat acts via regulation of *daf-28* ILP expression to regulate dauer formation independently of, or in parallel to, TORC2

Although *daf-7* and *daf-28* expression is downregulated in *rict-1* mutants, aberrantly increased dauer formation in *rict-1* mutants is restricted to 27°C. A change in growth temperature to from 25°C to 27°C is also sufficient to induce dauer formation at a low frequency in wild-type animals under otherwise similar cultivation conditions (27). To explore the mechanistic basis for enhanced dauer formation at 27°C, we quantified *daf-7* and *daf-28* expression levels across a range of cultivation temperatures. *daf-7*p::*gfp* expression was previously reported to be downregulated at 27°C (27). However, under our growth conditions and consistent with another report (62), we observed no obvious reduction in *daf-7*p::*gfp* expression, but instead found expression to be slightly increased in wild-type L1 larvae (F∽2.8, Df=1, P∽0.099; Figure 3A). This observation suggests that temperature-dependent downregulation of TGF-β transcription may not be the primary driver of dauer formation at 27°C.

**Figure 3.**
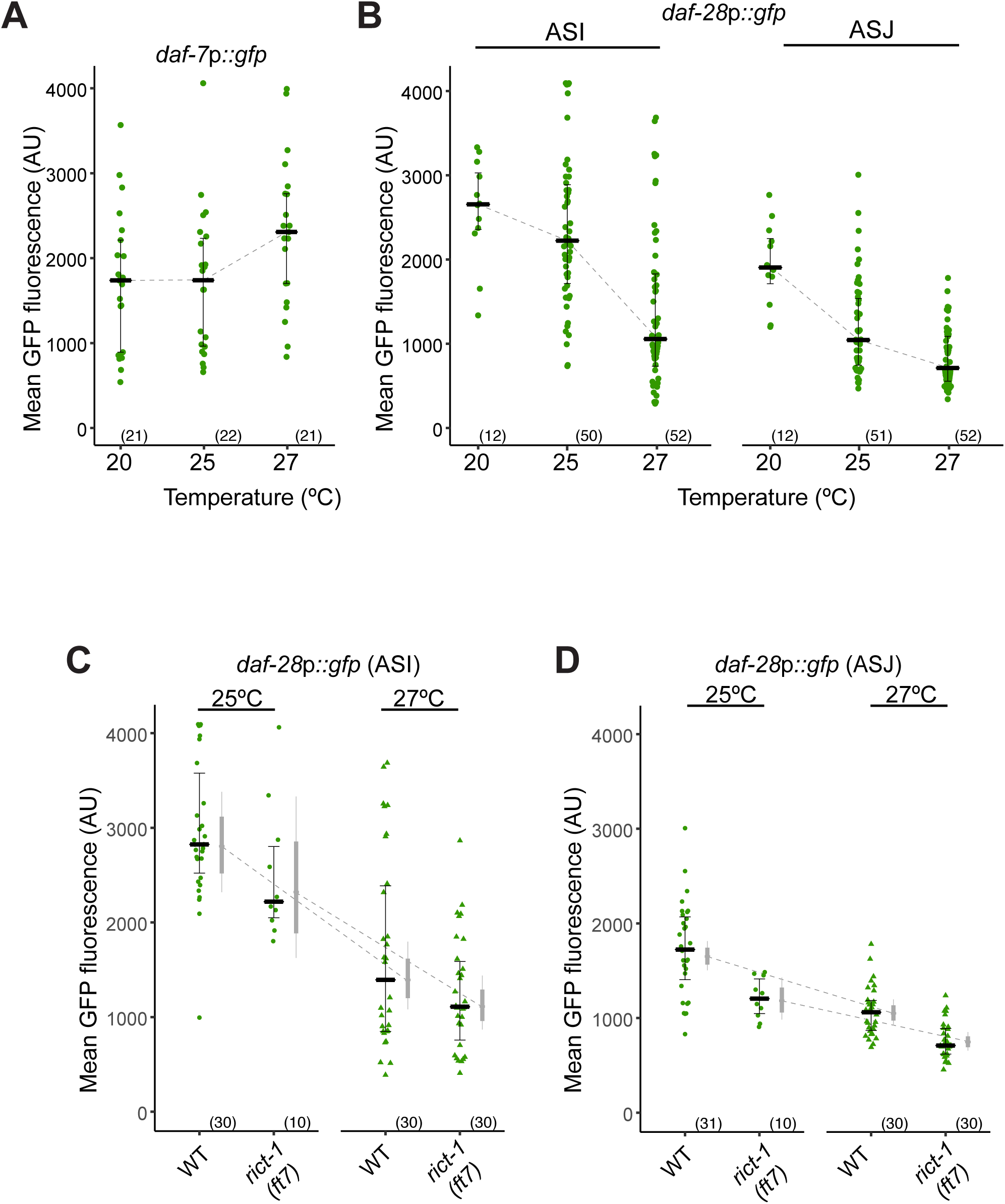
Parallel modulation of neuroendocrine signaling by heat and Rictor regulates dauer formation. **A-D)** Quantification of *daf-7*p::*gfp* and *daf-28*p::*gfp* fluorescence in the ASI, and ASI and ASJ neurons, respectively, in L1 larvae grown at the indicated temperatures. Each data point is the average GFP fluorescence of a single animal (2 neurons per animal); numbers in parentheses below indicate the total number of animals examined. Triangles in (C,D) indicate data that are repeated from Figure 2. Horizontal lines are the median; error bars are quartiles. Light gray thin and thick vertical bars at right of each scatter plot in (C,D) indicate Bayesian 95% and 75% credible intervals, respectively. Dashed lines indicate slope of median (A,B) or estimated mean (C,D) as a function of temperature.

In contrast, we found that *daf-28*p::*gfp* expression was strongly decreased in both the ASI and ASJ neurons as a function of temperature in wild-type L1 larvae (for ASI F∽24.1, Df=1, P < 0.001; for ASJ F∽49.0, Df =1, P < 0.001; Figure 3B). In particular, the expression of *daf-28*p::*gfp* in ASI decreased more steeply when growth temperatures were increased from 25°C to 27°C as compared to expression level changes from 20°C to 25°C (Figure 3B). *daf-28*p::*gfp* expression levels were decreased in *rict-1* mutants relative to those in wild-type larvae at all examined temperatures (Figure 3C,D). Nevertheless, these mutants exhibited a further reduction in *daf-28*p::*gfp* expression at higher temperatures (Figure 3C,D). These results suggest that decreased *daf-28* expression at 27°C drives dauer formation in both wild-type animals and in *rict-1* mutants, and that lower baseline expression levels of both *daf-28* and *daf-7* at 27°C amplifies this effect in *rict-1* mutants resulting in the Hid phenotype.

Dauer formation is increased upon the addition of exogenous pheromone (24, 25).At 25°C, pheromone promotes dauer formation via downregulation of *daf-7* and *daf-28* expression in ASI (28). We reasoned that the lower baseline levels of TGF-β and insulin signaling in *rict-1* mutants would also enhance pheromone-induced dauer formation in these animals at lower temperatures. Consistent with this hypothesis, we found that *rict-1* mutants exhibited increased dauer formation in response to the ascr#5 synthetic ascaroside (63) at 25°C (Figure S1B). Thus, lower levels of *daf-7* and *daf-28* expression in *rict-1* mutants sensitizes these animals to both temperature- and pheromone-induced dauer entry.

### *rict-1* mutants are insensitive to bacterial food quality

The feeding state of *C. elegans* larvae is reflected in decreased levels of both *daf-7* and *daf-28* signaling: downregulation of both pathways in food-restricted animals enhances dauer entry under dauer-inducing conditions at 25°C (28, 31, 36-38). Given the intestinal site of action of RICT-1 in the regulation of dauer formation, and the reduced expression of both neuroendocrine signaling pathways in *rict-1* mutants, a plausible explanation for these observations is that *rict-1* mutants exhibit defects in detecting and/or transmitting food information to regulate hormonal signaling.

We found that growth on live *E. coli* HB101 (high food quality) (41, 64) suppressed high temperature-induced dauer formation in wild-type animals as compared to growth on live *E. coli* OP50 (low food quality) (Figure 4A). Consistent with a role for food in regulating hormonal signaling, wild-type animals displayed increased *daf-28*p::*gfp* expression in the ASJ neurons when grown on HB101 compared to OP50 (Figure 4B). However, the Hid phenotype of *rict-1* mutants was unaffected by bacterial food quality (Figure 4A). Since the penetrance of the Hid phenotype of *rict-1* mutants is nearly 100% (Figure 4A), it is possible that we failed to observe an effect of the bacterial strain on dauer formation due to a ceiling effect. However, we found that expression of *daf-28*p::*gfp* was also not modulated by the identity of the bacterial strain in *rict-1* mutant animals (Figure 4B). We infer that Rictor is required to mediate food quality-dependent changes in *daf-28* ILP expression to regulate dauer formation.

**Figure 4.**
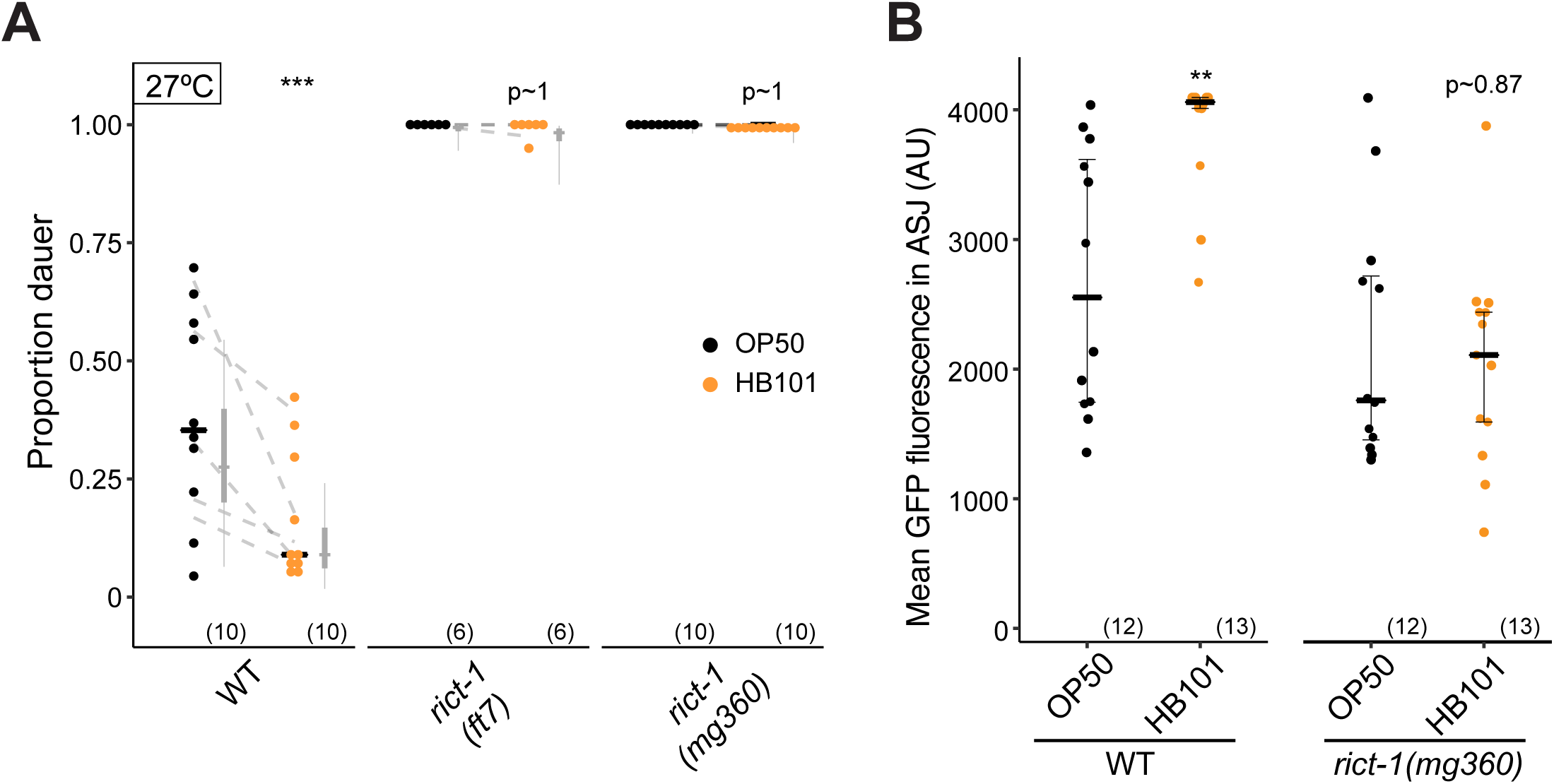
*rict-1* Rictor mutants are insensitive to food quality and feeding state. **A)** Dauers formed by wild-type and *rict-1* mutants on OP50 or HB101 at 27°C. Each black dot indicates the average number of dauers formed in a single assay. Horizontal black bar indicates median. Light gray thin and thick vertical bars at right indicate Bayesian 95% and 75% credible intervals, respectively. Numbers in parentheses below indicate the number of independent experiments with at least 19 animals scored in each. Dashed lines indicate mean change in dauer formation per experimental day. *P*-values are with respect to growth on OP50; *** - different from growth on OP50 at *P*<0.001 [two-factor ANOVA with F(5.20) with 2 Df, P=0.0092 for genotype*food interaction, Tukey-type multivariate-t post-hoc adjustment]. **B)** Quantification of *daf-28*p::*gfp* fluorescence in ASJ neurons of wild-type and *rict-1* mutant L2 larvae grown on OP50 and HB101 at 27°C. Each data point is the average GFP fluorescence of a single animal (two neurons per animal); numbers in parentheses below indicate the total number of neurons examined. Horizontal lines are the median; error bars are the quartiles. *P*-values are with respect to growth on OP50; ** - different from growth on OP50 at *P*<0.01 [two-factor ANOVA with F(7.76) with 1 Df, P=0.0077 for genotype*food interaction, Tukey-type multivariate-t post-hoc adjustment].

### Rictor acts in the intestine to regulate adult foraging behavior via PDF-1 signaling

We next investigated whether intestinal RICT-1 also plays a role in regulating food-dependent foraging behaviors in adult *C. elegans*. The low activity state (dwelling) is characterized by high angular velocity and reduced locomotion of animals on a bacterial lawn, whereas high activity exploratory behavior (roaming) is marked by reduced turning frequency and increased forward movement (40). Using a previously described assay (42) (Figure 5A), we found that as expected, wild-type animals fed *ad libitum* overnight did not roam extensively when placed on an OP50 lawn (Figure 5B). In contrast, fed *rict-1* mutants exhibited an approximately two-fold increase in exploratory behavior (Figure 5B). Automated videotracking of animal locomotion further confirmed that *rict-1* mutant adults spend a larger fraction of time roaming (Figure 5C-D). As in the case of dauer formation, expression of genomic *rict-1,* as well as expression of wild-type *rict-1* transgene in the intestine rescued the increased roaming behavior of *rict-1* mutants (Figure 5B). The roaming behavior of both wild-type and *rict-1* mutants was further enhanced by prolonged starvation (Figure S2A,S2B), indicating that *rict-1* mutants remain sensitive to food removal. We conclude that RICT-1/TORC2 acts in the intestine to modulate food-dependent adult behaviors.

**Figure 5.**
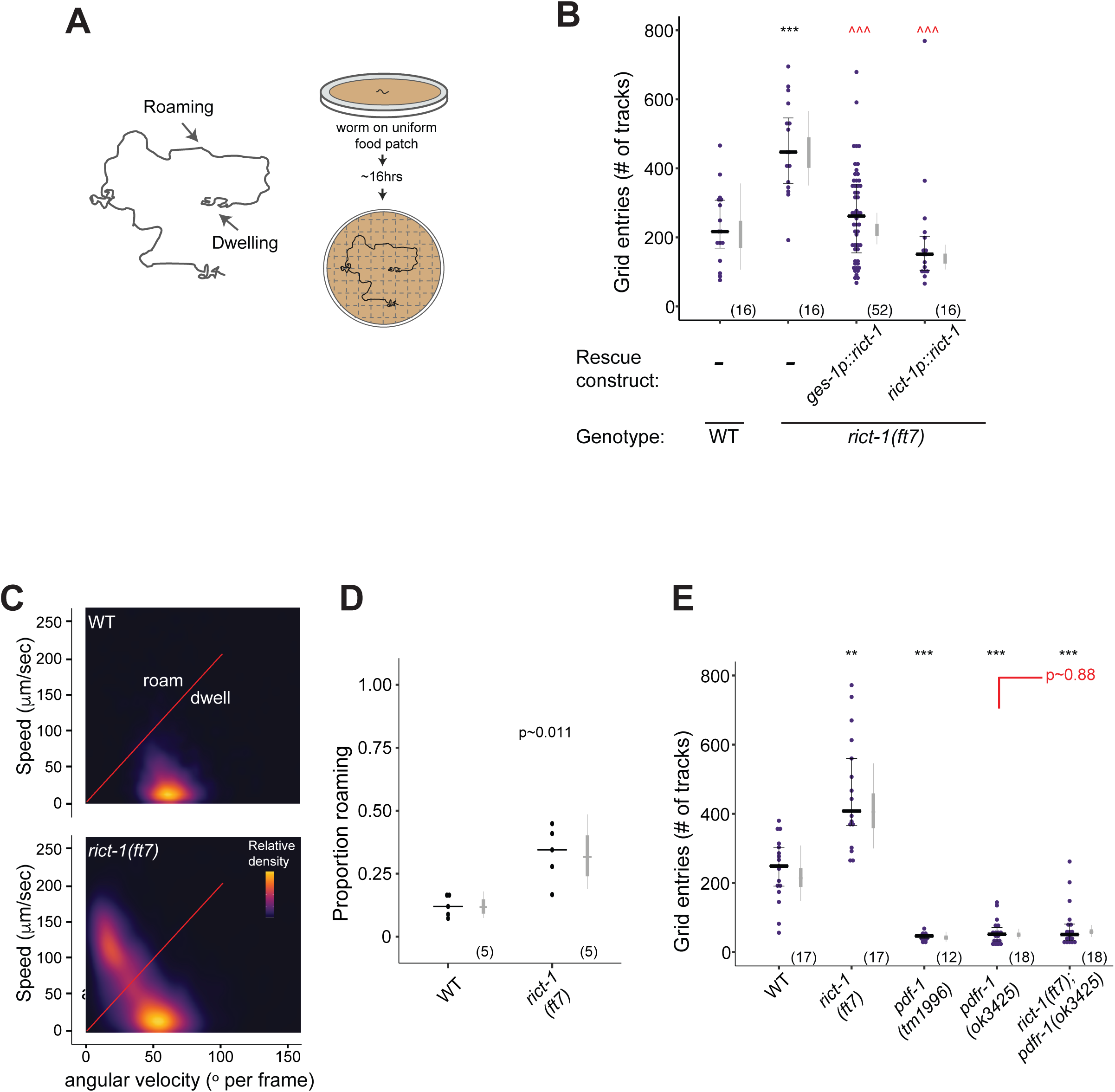
Intestinal RICT-1 regulates adult exploratory behaviors. **A)** Cartoon depicting foraging assay. Each plate contains a L4 hermaphrodite which is allowed to forage overnight on a uniform lawn of *E. coli* for ∽ 16hrs. Lines show tracks on the bacterial lawn. **B,E)** Exploratory behavior of indicated strains. Each purple dot represents data from a single worm. Median is indicated by a horizontal purple line; error bars are quartiles. Light gray thin and thick vertical bars at right indicate Bayesian 95% and 75% credible intervals, respectively. Numbers in parentheses below indicate the total number of animals examined during 3 independent days of assays. *P*-values shown are between indicated values; *** and ^^^ - different from WT and *rict-1* at *P*<0.001, respectively (ANOVA with Tukey-type multivariate-t post-hoc adjustment). *P*-values of differences in means relative to wild-type and corresponding mutant animals are indicated in black and red, respectively. **C)** Live tracking of foraging behavior. Shown is a representative assay. n > 20 animals per assay. Density plot shows mean speed and angular velocity from tracks binned over 10 sec intervals imaged at 3 frames/sec. Red line indicates delineation of roaming (upper region) and dwelling (lower region) behavior. **D)** Quantification of roaming and dwelling states. Dots show proportion of track time bins in which animals were roaming, with each dot reflecting one assay. Light gray thin and thick vertical bars at right indicate Bayesian 95% and 75% credible intervals, respectively. *P*-value shown is with respect to wild-type (Welch’s t-test).

Switching between the states of roaming and dwelling is in part regulated by hormonal and neuropeptide signaling. DAF-7 has previously been shown to modulate a behavioral switch between the fasted/fed state of quiescence and dwelling, but not the switch between roaming and dwelling (21, 43, 65, 66), suggesting that RICT-1 modulates adult foraging behaviors via alternate mechanisms. The PDF-1 neuropeptide has been shown to act via the PDFR-1 receptor to modulate exploratory behaviors; *pdf-1* and *pdfr-1* mutants display reduced roaming by altering the duration of roaming and dwelling states (42). To determine whether *rict-1* acts via regulation of PDFR-1 signaling, we examined the foraging phenotypes of *rict-1; pdfr-1* double mutants. We found that the exploratory behavior of *rict-1; pdfr-1* double mutants resembled those of *pdfr-1* single mutants alone, suggesting that *pdfr-1* is epistatic to *rict-1* for foraging phenotypes (Figure 5E). *pdfr-1* mutations did not suppress the Hid phenotype of *rict-1* mutants (Figure S2C), indicating that intestinal RICT-1 regulates dauer formation and adult foraging behaviors via distinct neuroendocrine signaling pathways.

### Dauer formation and adult exploratory behavioral phenotypes result from distinct genetic contributions

Since TORC2 regulates both dauer formation and adult exploratory behaviors, we investigated the extent to which these traits covary in the population. To address this issue, we examined high temperature-induced dauer formation and adult foraging behaviors in a panel of 22 strains, representing 19 unique isotype sets isolated from various latitudes (67). We found that high temperature-regulated dauer formation varied among these strains (Figure 6A) and was uncorrelated with isolation latitude and pheromone-driven dauer formation (Figure S3). We next analyzed the foraging behaviors of a subset of the wild isolates exhibiting the most extreme dauer formation phenotypes.With the exception of the CB4856 Hawaiian and PX178 strains that displayed increased exploratory behavior, the foraging behaviors of adult animals from examined strains were similar to those of the laboratory N2 strain (Figure 6B). Based on this limited dataset, we conclude that the genetic contributions to high temperature-induced dauer formation and foraging behavior are largely separable in the wild population.

**Figure 6.**
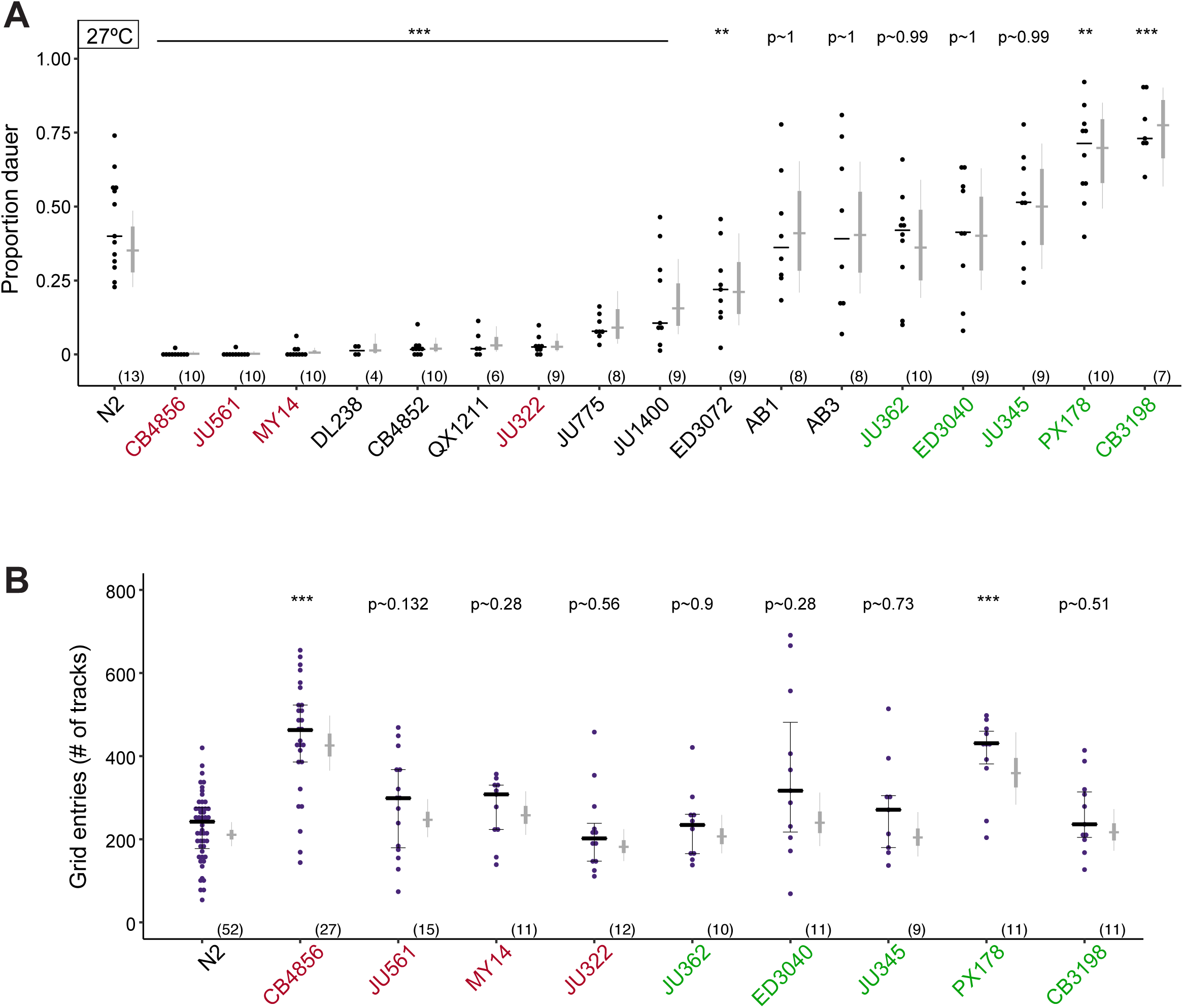
High temperature-induced dauer formation and adult exploratory behaviors do not covary. **A)** Dauers formed by the indicated *C. elegans* strains at 27°C. Strains also assayed for exploratory behavior in (B) are in red (weaker Hid phenotype than Bristol N2) and green (similar or stronger Hid phenotype than N2). Each black dot indicates the average number of dauers formed in a single assay. Horizontal black bar indicates median. Light gray thin and thick vertical bars at right indicate Bayesian 95% and 75% credible intervals, respectively. Numbers in parentheses below indicate the number of independent experiments with at least 16 animals each. All shown *P*-values are with respect to the Bristol N2 strain; ** and *** - different from the Bristol N2 strain at *P*<0.01 and *P*<0.001, respectively (ANOVA with Dunnett-type multivariate-t post-hoc adjustment). **B)** Exploratory behavior of indicated strains. Strains in red and green exhibit weaker or similar/stronger Hid phenotypes than N2, respectively (A). Each purple dot is data from a single animal. Median is indicated by a black horizontal line; error bars are quartiles. Light gray thin and thick vertical bars at right indicate Bayesian 95% and 75% credible intervals, respectively. Numbers in parentheses below indicate the total number of animals examined in at least 3 independent assay days. *P*-values shown are with respect to N2; *** - different from N2 at *P*<0.001 (ANOVA with Dunnett-type multivariate-t post-hoc adjustment).

### A single locus largely accounts for inter-strain variation in high temperature-induced dauer formation in N2 and CB4856

In contrast to both N2 and *rict-1* mutants, the CB4856 strain is strongly defective in dauer formation at 27°C (Figure 6A). However, similar to *rict-1* mutants, CB4856 animals exhibit increased roaming behavior on food (Figure 6B). To identify additional loci that contribute to dauer formation and/or roaming behavior, we sought to identify causative loci for these differences between N2 and CB4856. We first utilized a collection of chromosome substitution strains (CSSs) in which a single CB4856 chromosome has been introgressed into an otherwise N2 genetic background (68) to initially map the loci of major effect in dauer formation. We found strong statistical support for one or more CB4856 loci on chromosome II contributing to a negative effect on dauer formation (LRT=16.17, Df=1, p=5.8x10^-5^), while introgression of CB4856 chromosome X (LRT=3.46, Df=1, p=0.054) and IV (LRT=2.75, Df=1, p=0.087) may contribute minor effects (Figure 7A). The CB4856 loci on chromosome II may generally affect entry into the dauer stage, or could specifically contribute to high temperature-induced dauer formation. We found that although the CSSII strain exhibited strong defects in high temperature-induced dauer formation, these animals retained the ability to form dauers in response to the ascr#5 pheromone (Figure S4A). This result suggests that CB4856 variants on chromosome II specifically modulate dauer formation in response to high temperature.

**Figure 7.**
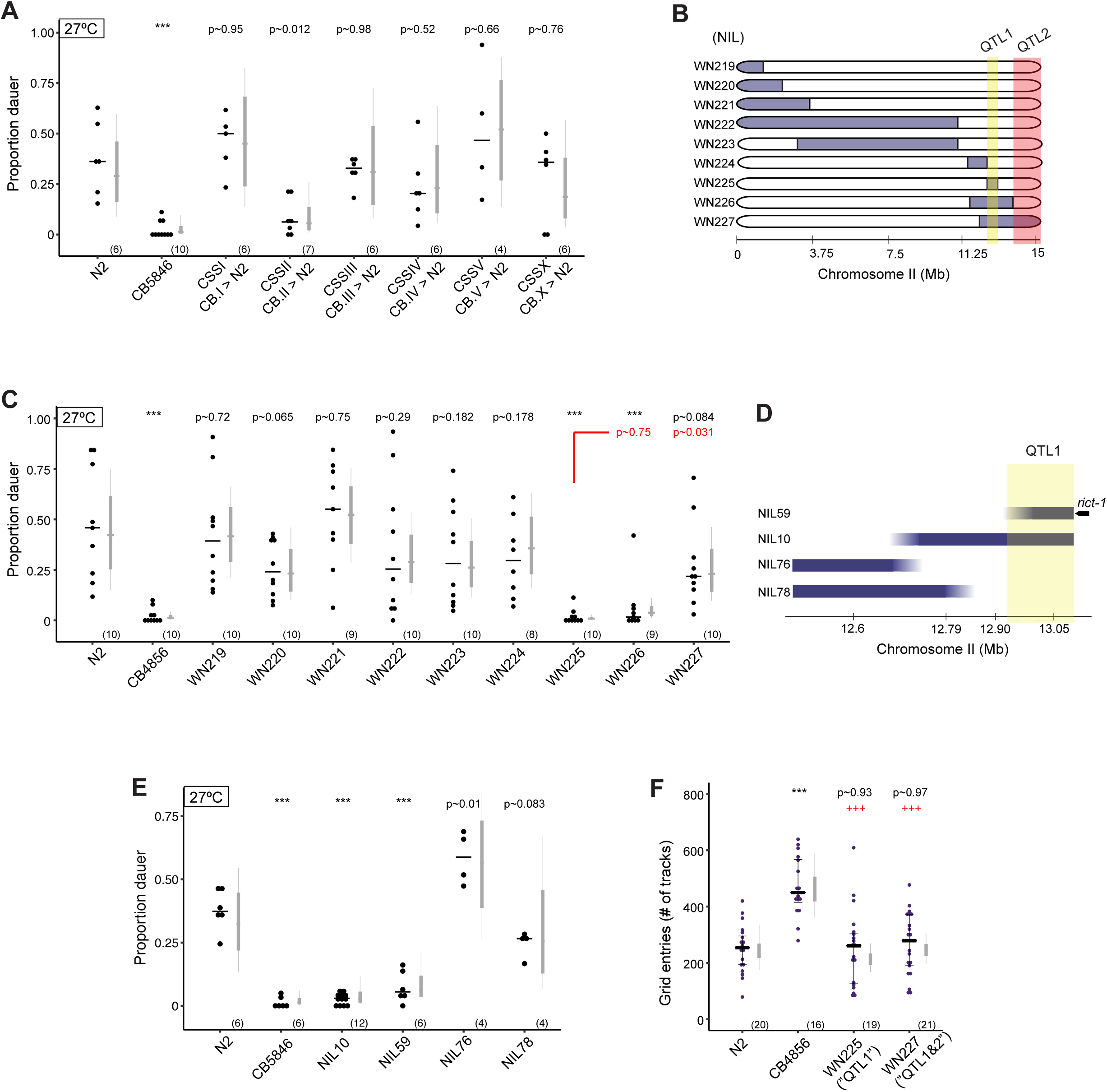
Identification of a locus contributing to variation in high temperature-induced dauer formation between N2 and CB4856. **A,C,E)** Dauers formed by chromosome-substituted strains (A) and NILs (C,E) at 27°C. Each dot indicates the proportion of dauers formed in a single assay. Horizontal bar indicates median. Light gray thin and thick vertical bars at right indicate Bayesian 95% and 75% credible intervals, respectively. Numbers in parentheses below indicate the number of independent assays with at least 11 animals each. Shown *P*-values are with respect to the Bristol N2 strain unless indicated otherwise; *** - different from Bristol N2 at *P*<0.001 (ANOVA with Dunnett-type multivariate-t post-hoc adjustment for A, and Tukey-type multivariate-t adjustment for C and E). **B,D)** Cartoons depicting CB4856 introgression breakpoint in NIL strains. Segments in blue indicate regions of CB4856 DNA in an otherwise Bristol N2 genetic background. NILs in B have been described (69). Yellow shaded region indicates likely position of QTL1 and orange indicates probable location of QTL2. Horizontal axis indicates physical position in megabases (Mb). **F)** Exploratory behavior of indicated strains. Each purple dot is data from a single animal.Median is indicated by a horizontal line; error bars are quartiles. Light gray thin and thick vertical bars at right indicate Bayesian 95% and 75% credible intervals, respectively.Numbers in parentheses below indicate the total number of animals examined in 3 independent assay days. *P*-values shown are with respect to N2; ^+++^ - different from CB4856 at *P*<0.001 (ANOVA with Tukey-type multivariate-t post-hoc adjustments).

To identify the loci on chromosome II that may account for the dauer formation effect, we next analyzed a collection of near-isogenic lines (NILs) that carry CB4856 chromosomal segments introgressed into the N2 genetic background, and that cover most of chromosome II (69) (Figure 7B). Through analysis of dauer formation at 27°C, we identified a single locus that largely phenocopies the CB4856 strain in this assay (Figure 7C). Using the minimal locus of largest effect, we constructed sub-NILs to further refine this interval (Figure 7D). We identified an interval of approximately 100 kb on chromosome II (‘QTL1’) that nearly fully recapitulates the CB4856 effect on dauer formation when introgressed into the N2 background (Figure 7E). Accounting for the phenotypic effects of QTL1, we also found evidence for an additional QTL (“QTL2”), which appears to act in opposition to QTL1 to regulate dauer formation (Figure 7B,C).Although QTL1 maps in proximity of the *rict-1* gene, analysis of the breakpoints of the smallest interval for QTL1 showed that it does not include the coding region of *rict-1* (Figure 7D, Figure S4B) (70). Consistent with this observation, although the *rict-1* gene in CB4856 contains two missense polymorphisms relative to N2 sequences, conversion of residues in the N2 strain to the CB4856 variants using CRISPR-mediated gene editing did not alter dauer formation at high temperatures (Figure S4C). These results suggest that one or more variants in the QTL1 and QTL2 intervals account for the CB4856 dauer formation phenotype.

To determine whether similar to *rict-1* mutants, these QTLs affect both dauer formation and adult exploratory behaviors, we analyzed foraging behavior in these NIL strains. We found no significant change in foraging behavior in these NILs (Figure 7F) suggesting that additional QTLs contribute to the increased exploratory behavior of the CB4856 strain. Thus, consistent with our analysis of wild isolates, foraging behaviors and dauer formation appear to be largely genetically separable.

## DISCUSSION

In this study, we show that food-dependent regulation of a developmental decision and an adult behavior is mediated in part by Rictor/TORC2 in the intestine. Intestinal TORC2 transmits internal metabolic state information to regulate neuroendocrine pathways, establishing this complex as a key player in gut-to-brain communication (Figure 8). Our results indicate that multimodal sensory inputs and internal state information converge to regulate neurohormonal signaling, thereby enabling animals to integrate diverse environmental cues in the context of their experience and current conditions.

**Figure 8.**
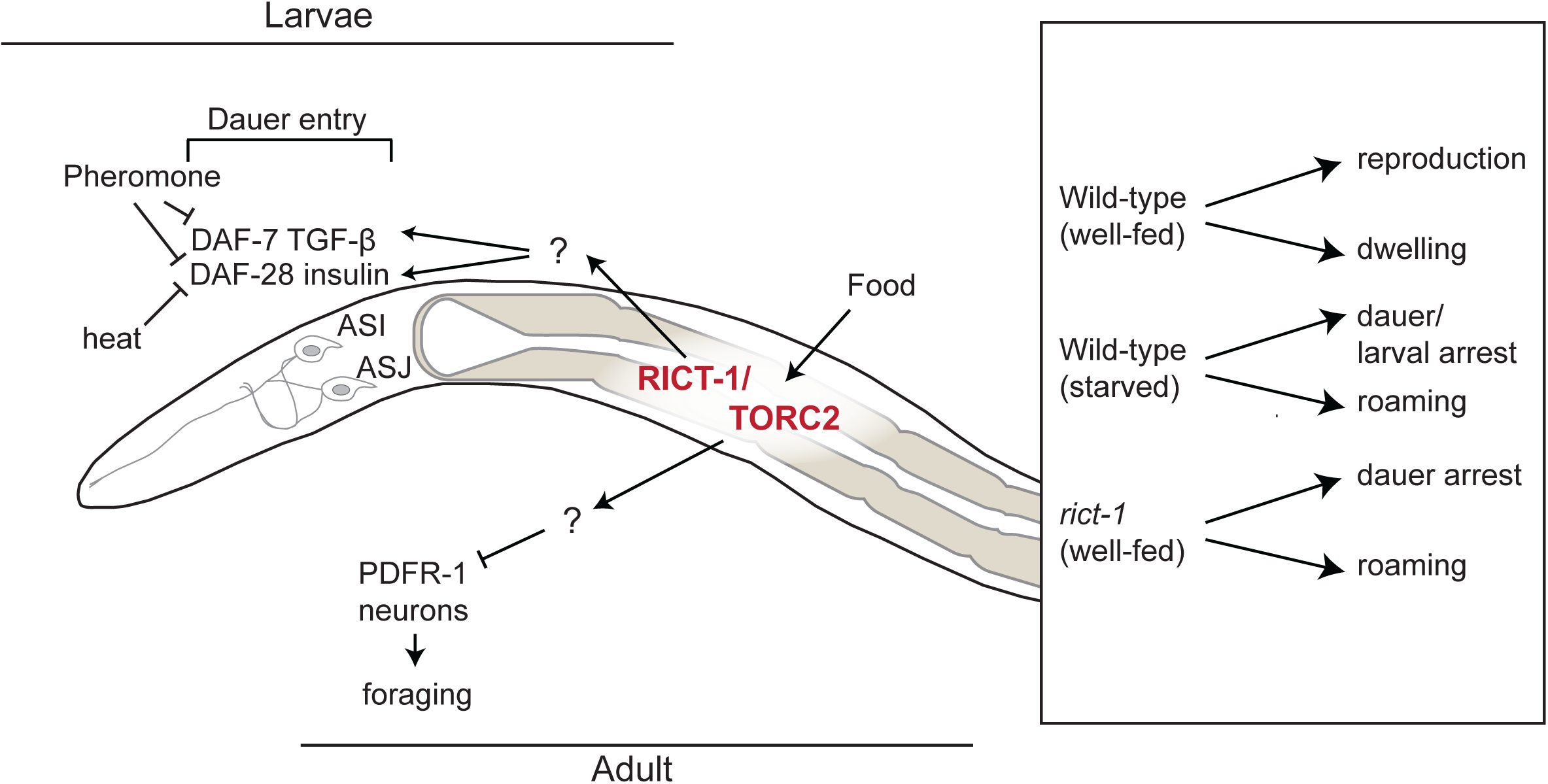
Model for weighting of environmental cues in the regulation of dauer development and behavior. In larvae, pheromone primarily downregulates expression of *daf-7* and *daf-28* in ASI, whereas heat downregulates *daf-28* expression in both ASI and ASJ to promote dauer formation. TORC2 signaling acts in the gut in a diet-dependent manner to target both *daf-7* and *daf-28* expression, thereby allowing hierarchical integration of internal metabolic state signals with external cues. TORC2 signaling in the gut also acts via PDFR-1 to modulate adult exploratory behaviors.

### Hierarchical control of dauer formation

TGF-β and insulin signaling act in parallel to regulate dauer formation (30). Our results together with previously published work indicate that expression of *daf-7* TGF-β and *daf-28* ILP (as well as other ILP) genes is regulated combinatorially by multiple sensory inputs (28, 31-33, 35-37). While expression of *daf-7* in ASI is strongly downregulated by pheromone (28, 32-34), crude pheromone extracts downregulate *daf-28* expression primarily in ASI but not in ASJ (28, 31) (Figure 8). Conversely, we find that *daf-7* expression is largely unaffected by high temperature, whereas *daf-28* expression in both ASI and ASJ is strongly reduced upon a temperature shift to 27°C (Figure 8; this work). In contrast to the effects of temperature and pheromone, starvation robustly downregulates expression of both *daf-7* and *daf-28* in larvae (28, 31-33, 36-38). Consequently, low food quantity and/or quality is a potent enhancer of both pheromone-as well as temperature-induced dauer formation (20, 28) (this work). These observations imply a hierarchical control of dauer formation; high food quality/availability can override the dauer-instructive cues of pheromone and temperature at the level of regulation of neuroendocrine gene expression to promote reproductive development (Figure 8). A similar hierarchical set of signals has been shown to modulate *daf-7* expression in ASJ in males in response to multiple internal and external inputs to control adult male exploratory behavior (71).

The decision to exhibit one of alternate polyphenic traits requires weighing the predictive value of an uncertain environment against the cost of entering into the inappropriate stage (72, 73). In natural settings, pheromone levels and temperature are likely to fluctuate over the time period during which the decision to commit to a specific developmental pathway is made (26, 74). However, metabolic signaling of nutrient status originating from the gut may be expected to vary on longer timescales than externally derived cues, and may thus have a higher predictive value for current and future environmental conditions. By regulating both the TGF-β and insulin pathways, internal feeding state signals may, therefore, strengthen or weaken the salience of pheromone and temperature signals, and appropriately bias larval developmental decisions.

### A role for intestinal TORC2 in nutrient sensing

Unlike mTORC1, which has been definitively shown to be activated by stress, amino acids, energy status and oxygen levels, a role for TORC2 in direct nutrient sensing is less well-established (47, 48, 75). However, recent work suggests that TORC2 may respond to metabolic signals, such as those involved in glutamine metabolism (76). In the context of dauer development and adult foraging behaviors, *rict-1* mutants exhibit phenotypes that qualitatively mimic the effects of nutrient depletion in wild-type animals. For instance, *daf-7* and *daf-28* expression is downregulated in both starved wild-type and fed *rict-1* mutants, and the exploratory behaviors of *rict-1* mutants resemble those of food-deprived wild-type animals. These phenotypes are unlikely to arise simply from defective feeding, because pharyngeal pumping rates were shown to be either unchanged or increased in *rict-*1 mutants (53, 55). Taken together with the observation that expression of *rict-1* in the intestine is sufficient to rescue its dauer development and foraging phenotypes, we hypothesize that TORC2 plays a role in sensing or metabolism of nutrients in the *C. elegans* intestine. However, while Rictor activity in the gut can regulate traits such as growth rate and fat storage, TORC2 also acts in neurons to regulate lifespan (11, 54). Moreover, TORC2 has recently been reported to act in ASI to regulate hunger-induced changes in aversive behaviors (77). Thus, it remains formally possible that TORC2 also acts in a subset of neurons to regulate dauer development and adult foraging.

What signals from bacteria might activate TORC2 in the gut? The OP50 B strain is considered to be less nutrient-rich than the K12-derived strains HB101 or HT115, with the latter containing higher carbohydrate levels as well as differential profiles of amino and fatty acids (78, 79). Additionally, although differences in the levels of L-Trypophan (Trp) metabolites in B and K strains may contribute to food-dependent differences in reproductive output (80), Trp levels do not appear to account for food quality effects on longevity in *rict-1* mutants (54). Thus, the relevant differences in nutrient profiles of these food sources are likely to be complex (81). In the future, identification of the metabolites produced by these bacteria should yield insights into the differential regulation of phenotypes by different bacterial strains in *C. elegans*, as well as the roles of these nutrients in activating TORC2 signaling.

### Divergence in TORC2-regulated gut-to-brain signaling pathways

Although TORC2 signaling in the gut targets *daf-7* and *daf-28* in sensory neurons to regulate dauer formation, the neuronal targets of TORC2 in the regulation of adult foraging behaviors are likely to be distinct. This conclusion is based on the observation that mutations in *pdfr-1* suppress the roaming but not Hid phenotype of *rict-1* mutants.The TORC2-dependent pathways that transmit nutrient information directly or indirectly between the gut and sensory neurons in the head in either the larva or in the adult are currently unknown. Multiple members of the large ILP family are expressed in the gut, and expression and function of these genes in the intestine and other tissue types is regulated in a spatiotemporally complex manner in response to environmental signals and internal conditions (35, 37, 38, 82-84). In addition to its other well-described transcriptional outputs, TORC2 has been suggested to regulate histone modification via the dosage compensation complex in *C. elegans* (85). It is possible that TORC2 regulates the expression of different ILP genes in the intestine in response to distinct energetic conditions, and that this ILP ‘code’ and network (82) relays internal state information from the gut to the nervous system [and possibly also in the reverse direction (84)] to modulate developmental and behavioral outcomes (12, 38, 82, 86) (Figure 8).

The complexity and divergence of pathways regulating sensory signal-driven dauer formation and foraging is reflected in our observations that these traits do not cosegregate in wild strains. We also identified two QTLs that act antagonistically to regulate high-temperature-driven dauer formation, but which do not affect pheromone-dependent dauer formation or foraging decisions, suggesting that the genetic architectures underlying these phenotypes are distinct. Rictor/TORC2 may represent a common genetic node in these networks that include diverse sensory input pathways, as well as diverse sets of signals originating from the gut to regulate multiple phenotypic outcomes. Further characterization of these pathways both in lab-derived and in wild strains may allow us to determine the mechanisms by which TORC2 is activated, and how this conserved protein complex mediates diverse downstream effects as a function of nutrient status.

## MATERIALS AND METHODS

### Strains

All strains were maintained on nematode growth medium (NGM) at 20°C using *E. coli* OP50 as a food source. Plasmids were injected at 5 ng/μl together with the *unc-122*p::*gfp* coinjection marker at 30 ng/μl to generate transgenic strains. At least two independent lines were examined for each transgene with the exception of *ges-1*p-driven constructs. Wild isolate strains were acquired from the *Caenorhabditis* Genetics Center (CGC, University of Minnesota). *E. coli* strains were streaked from glycerol stocks prior to use and grown to saturation in LB media at 37°C. Strains used in this study are described in Table S1.

### Generation of near-isogenic lines (NILs)

Construction of NIL strains has been previously described (69). Standard recombination techniques were used to generate sub-NILs. In brief, N2 males were crossed to WN225. F2 progeny derived from the F1 heterozygous progeny of this cross were screened for the presence of SNPs at 12.63 Mb (WBvar00001258, G>T in CB4856) using primers 12.63S 5’-TTGAACGTTCAAAAACTCCGTATCACG-3’ and 12.63AS5’-TGAAACCGTATTGCCATTGCATGC-3’ and at 12.991 Mb (WBvar00240486, C>T in CB4856) using primers 12.99S 5’-TTAGGAAAGTTGTGTCCACCTGGCGCGTGC-3’and12.99AS5’-CAAATTCCACTACGGGGGCACTGTCAATCAATTAGGCC-3’.Breakpoints of sub-NILs were identified by sequencing. The smallest sub-NIL containing QTL1 (NIL59) had a left breakpoint between 12.94 Mb (WBVar01380590) and 12.99 Mb, and a right breakpoint between 13.09 Mb (WBVar01380828) and 13.10 Mb(WBVar00003943). The resulting interval between 95-154 kb contains 20-29 genes, and 298-401 annotated variants, of which 24-34 are predicted non-synonymous protein-coding changes (70).

### Molecular biology

cDNA sequences for *rict-1b, sgk-1b* and *akt-1b* were amplified via PCR from a mixed stage cDNA pool and confirmed by sequencing. All promoter sequences were cloned from genomic DNA. The following promoters were used in this study: *ges-1p* (strong intestine, weak in non-neuronal head cells, 2.1 kb), *elt-2*p (strong intestine, 3.3 kb), *ifb-2*p (strong intestine, 3 kb), *gpa-4*p (ASI, 2.6 kb). Rescue constructs were expressed along with a bicistronic *SL2*::*mCherry*::*unc-54* 3’UTR cassette. The expression pattern of intestinal promoters was confirmed using GFP reporters. *srg-47*p::*daf-7* (ASI), *srg-47*p::*daf-28* (ASI) and *trx-1*p::*daf-28* (ASJ) have been described previously (28).Vector maps are available on GitHub (https://github.com/mikeod38/dauergut.git). CB4846 *rict-1* SNPs were introduced into the N2 strain via pDD162-plasmid-based CRISPR/Cas9 gene conversion using the *dpy-10* co-CRISPR technique (87, 88). sgRNA sequences were cloned via PCR mutagenesis using the pU6-*klp-12* sgRNA as a template (89). Cas9 plasmid pDD162 was injected at 50 ng/μl and sgRNA plasmids were injected at 25 ng/μl. Donor oligos were injected at 500 nM. Donor oligo sequences for *rict-1* were: WBVar01249304(D1462G)5’-TTTCCAGCCAAACCGAAATCGGATCCGATCTTTTCATTCCACGAGAACGGCGACTCTGCAGGCGTCGAGGATCGTGGAGCCCGAACCGGACACGC-3’,and WBVar00004038(N1174K)5’-AGGAAGATTACTATTCAGAGTCATCGGAAATCTTCGATGGTTGATGTGAGTTAGGGCAAA-3’.

### Dauer formation assays

All examined strains were cultured for at least 3 generations without being allowed to starve prior to use in dauer assays. Dauer formation assays were performed essentially as described (90). In brief, 5 growth-synchronized young adult animals grown on OP50 were placed on dauer assay agar plates (90) with (for pheromone-induced dauer formation assays) or without (for heat-induced dauer formation assays) ascr#5 pheromone and live *E. coli. E. coli* was spun down from a saturated overnight culture and resuspended at 16 μg/μl in S-basal medium containing 5 μg/ml cholesterol. 160 μg or 80 μg of this suspension was plated per assay plate to assess high-temperature or pheromone-driven dauer formation, respectively, and allowed to dry. Worms were allowed to lay between 50-75 eggs per plate at room temperature. Adults were then removed and plates were placed at either 27°C or 25°C to assay temperature or pheromone-driven dauer formation, respectively. Dauers were identified visually using a dissection microscope after approximately 60 hrs to allow slower-growing strains to develop. Under these conditions we did not observe substantial dauer exit in the N2 strain as reported previously using different assay conditions (27).

### Microscopy

All fluorescence microscopy was performed using animals anesthetized with 100 mM levamisole (Sigma Aldrich). Animals were imaged on 2% agarose pads using an upright Zeiss Axio Imager with a 63X oil immersion objective. For larval imaging, animals were grown under identical conditions as those used for dauer formation assays with the exception that 160 μg of OP50 or HB101 was used regardless of assay temperature. All images were collected in *z*-stacks of 0.5 μm through the head of the animal. Quantification was performed using ImageJ (NIH). Fluorescence was quantified by identifying the focal plane in which most of the cell soma was visible, followed by manually drawing an ROI around the soma. Mean pixel intensity was recorded for each neuron pair per animal and the average of fluorescence in each animal is shown.

### Fluorescent *in situ* hybridization

Fluorescent *in situ* hybridization of *daf-7* was performed as described (91, 92).Animals were grown to the L1 molt stage and fixed in 3.7% formaldehyde for 30 minutes. Animals were transferred to 70% EtOH after PBS washes and were stored at 4°C for less than one week. Hybridization with probes was performed at 30°C. *z*-stacks were acquired using an upright Zeiss Axio Imager with a 63X oil immersion objective at 0.5 μm intervals. The ASI soma were identified by position. For quantification, sum projections from 4 focal sections covering 2 μm were analyzed using ImageJ (NIH). Mean pixel intensity was calculated using a manual ROI around each cell soma and background fluorescence was subtracted from this value to obtain normalized fluorescence.

### Exploration Assays

Foraging behavior assays were performed as described (42) with minor modifications. 6 cm NGM agar plates were uniformly seeded with 200 μl of a saturated culture of OP50 and allowed to dry overnight. Individual L4 animals were picked to these plates and transferred to 20°C for ∽14 hrs. Animals were then removed, and plates were photographed using a PixelLink D722MU-T camera mounted to a Nikon SMZ745 dissection scope with a 0.5X lens. Images were analyzed using a custom ImageJ (NIH) macro to superimpose a grid containing 188 squares. The number of times the worm had entered a square was manually counted based on the presence of tracks. An upper limit of 10-grid entries per box was set due to our inability to unambiguously identify additional tracks. This scoring method generated data that is approximately Poisson-distributed (see Statistical Analyses). Transgenic and mutant strains were assayed in parallel with control animals. For each genotype, a minimum of five animals was tested per day on at least three independent days.

### Live tracking of roaming/dwelling behavior

Live tracking and analyses of foraging behavior analysis was performed as described (42, 93) with modifications. 3.5 cm NGM plates were uniformly seeded with 200 μl of a saturated culture of OP50 and allowed to dry overnight. Prior to recording, 2 0 1d old adult animals were picked to a small spot on the edge of each assay plate, plates were placed at 20°C, and worms were allowed to move freely for 20 minutes.Temperature was maintained in an air flow-limited cabinet using a custom-built Peltier temperature control stage and plates were imaged using red LED circumferential illumination. Animals were recorded for 90 min at three frames per second using a PixeLink D722MU-T camera and a custom Matlab script. For off-food assays, worms were allowed to crawl freely on 3.5 cm NGM plates containing no food for 2 hours prior to assaying on food-free NGM plates.

Worm position (center of mass), speed, and direction were analyzed using WormLab software (MBF Bioscience, VT). Within WormLab, a Kalman filter was applied to the tracking data to minimize lateral sinusoidal translation of worm position. Data were analyzed using a custom R script to plot animal speed versus angular velocity. For angular velocity calculations, the position change over three frames at each time point was quantified. We identified two vectors: v1.2 representing the change in position from frame 1 to frame 2, and v2.3, representing the change in position from frame 2 to 3. The angle change per frame was defined as the dot product of these two vectors, divided by the scalar products of the norm of these two vectors. Angular velocity is expressed as degrees per frame.

For quantification of roaming and dwelling proportions, we visually identified clusters of data that were divided by a line with a slope of 2 on the speed/angular velocity scale. Points lying above and below that line were considered roaming and dwelling, respectively. For all conditions, at least 3 assays with two genotypes were performed in parallel on at least two independent days.

### Statistical Analyses

All statistical analyses were performed in R. Raw data as well as all code to replicate analysis are available on GitHub (https://github.com/mikeod38/dauergut.git).

*Dauer assays:* Dauer formation assays essentially consist of clustered binomial data, in which each assay plate comprises a cluster, with each animal representing an observation. The variability among clusters results in higher variance than what is expected for binomial data, requiring the application of more sophisticated statistical analyses.

A binomial generalized linear mixed effects model (GLMM) using the “lme4” R package was applied to estimate sources of variance in a large dataset of wild-type dauer formation at 27°C (n = 130 assays over 65 days). At least three different sources of variance were hypothesized: 1) day-to-day variance (sD), 2) plate-to-plate variance (sP), and 3) variance due to strain culture history (sG) which might be expected to affect strains on a given day in different directions. Modeling these variance terms as random effects in the GLMM fit suggested that the estimated magnitude of sD was greater than the estimate of sP, in agreement with observations of dauer formation. sG could not be directly estimated due to the absence of parallel, independently cultured wild-type samples for these data; however, sG is retained in these analyses.

To compare frequentist and Bayesian approaches in the statistical analyses of dauer formation assays, 1000 datasets were simulated under conditions of low and high variance (sD, sP and sG), and with different means and sample sizes. Two different scenarios were modeled: a) a comparison of two genotypes to a common control in which the true mean of all samples were equal, and b) comparison of these two genotypes in which one group differed from the control by a magnitude of 1 on the log-odds (logit) scale. Models also included the effects of including unbalanced data such as when all groups were not equally represented on each day of analysis, as well as estimations of the effects of systematic bias in the control samples, i.e. when a subset of the observation days had correlated mean values due to the effect of sD. For each comparison group, 6 data points (assay plates) sampled over 3 days were simulated. All simulations were performed in R.

Type-I and type-II error rates (Table S2) were modeled using four statistical methods: 1) multiple t-tests with Welch’s correction, 2) ANOVA, 3) binomial GLMM, and 4) Bayesian binomial GLMM, using the “rstantarm” R package with default, uninformative priors. For t-test and ANOVA, the response variable was the proportion of dauer formation, while the binomial response (dauer, non-dauer) of each animal was used for the GLMM methods. Random intercepts were fitted for sD, sP, and sG in GLMM, but were omitted for the t-test and ANOVA. The Bayesian binomial GLMM approach produced the lowest Type 1 error rates under conditions of high sD and sG, as well as with unbalanced data (Table S2), characteristics of many of the dauer assays reported here. For comparison, all figures are shown with *P*-values derived from ANOVA, as well as parameter estimates and credible intervals from a Bayesian GLMM.

*Roaming/Dwelling assays:* For grid-entry data, a plot of residuals from a linear model showed variance increasing with mean. It was hypothesized that the count data in these assays likely derives from a Poisson process. Thus, all data were analyzed using ANOVA and Bayesian GLMM with a Poisson link function and using default priors.Bayesian GLMM included a random effect term for “date” to account for sD and “plate” to account for sP.

*GFP fluorescence quantification*: A plot of residuals from a linear model showed variance increasing with mean, which was corrected using log10 transformation. Data were analyzed using ANOVA and a Bayesian LMM with random effects terms for “date” to account for sD and “plate” to account for sP.

## ACKNOWLEDGEMENTS

We are grateful to the *Caenorhabditis* Genetics Center, Dominique Glauser and Miriam Goodman for providing strains, Rebecca Butcher for the gift of ascr#5, Dennis Kim for providing *daf-7* FISH probes, Michael Ailion for advice, and Kyuhyung Kim, Scott Neal, Inna Nechipurenko and the Sengupta lab for critical comments on the manuscript. This work was supported in part by the NSF (NSF IOS 1655118 and NSF IOS 1256488 – P.S.), the NIH (R35 GM22463 – P.S., F32 DC013711 and T32 NS007292 – M.O’D.) and the Human Frontiers Science Program (RGP0028-2014 – J.K.).

## SUPPLEMENTAL FIGURE LEGENDS

**Figure S1.** Mutations in *rict-1* modulate dauer formation via downregulation of neuroendocrine signaling.

**A)** Dauers formed by animals of the indicated genotypes at 27°C. Each dot indicates the proportion of dauers formed in a single assay. Horizontal bar indicates median. Light gray thin and thick vertical bars at right indicate Bayesian 95% and 75% credible intervals, respectively. Numbers in parentheses below indicate the number of independent experiments with at least 26 animals each. *** - different from wild-type at *P*<0.001, ^^^-different from *rict-1(mg360)* at *P*<0.001 (ANOVA with Tukey-type multivariate-t post-hoc adjustments).

**B)** Dauers formed by wild-type (black) and *rict-1(ft7)* (blue) animals at 25°C in the presence of the indicated concentrations of ascr#5. Each dot indicates the proportion of dauers formed in a single assay. Horizontal bar indicates median. Numbers in parentheses below indicate the number of independent experiments with at least 27 animals each. Lines indicate predictions from GLM fit, corresponding to an odds ratio of ∽21.3 for *rict-1* across this range of ascr#5 concentrations. For genotype, Wald X^2^ = 45.3, Df =1, p < 0.0001; for pheromone, Wald X^2^ = 24.9, Df =1, p < 0.0001 with GLMM fit.

**Figure S2.** RICT-1 regulates dauer formation and adult exploratory behaviors via distinct hormonal signaling pathways.

**A)** Live tracking of foraging behavior in starved worms. Shown is a representative assay. n > 20 animals per assay. Density plot shows mean speed and angular velocity from tracks binned over 10 sec intervals imaged at 3 frames/sec. Red line indicates delineation of roaming and dwelling behavior.

**B)** Quantification of roaming and dwelling states. Dots show proportion of track time bins in which animals were roaming, with each dot reflecting one assay. Light gray thin and thick vertical bars at right indicate Bayesian 95% and 75% credible intervals, respectively. *P*-value shown is with respect to wild-type (Welch’s t-test).

**C)** *pdfr-1* mutations do not suppress *rict-1* dauer formation phenotypes. Each dot indicates the average number of dauers formed in a single assay. Horizontal bar indicates median. Light gray thin and thick vertical bars at right indicate Bayesian 95% and 75% credible intervals, respectively. Numbers in parentheses below indicate the number of independent assays with at least 27 animals each. *P*-value shown in comparison to *rict-1* mutants.

**Figure S3.** Isolation latitude of *C. elegans* strains is not correlated with high temperature-or pheromone-induced dauer formation.

Shown are the proportion of dauers formed at 27°C and in the presence of 6 μm ascr#5 pheromone by *C. elegans* strains isolated from different latitudes. Assays were performed on live OP50. For ascr#5 data (A-axis), each data point is the average of at least 3 independent assays of at least 23 animals each. 27°C data is repeated from Figure 6A. Error bars are the SEM. Line and shaded region indicate linear regression fit using mean dauer formation values for each strain.

**Figure S4.** Polymorphisms in *rict-1* coding sequences do not underlie the dauer formation phenotype of CB4856.

**A)** Pheromone-induced dauer formation is not reduced in the CSSII strain. Each dot indicates the average number of dauers formed in a single assay. Horizontal bar indicates median. Numbers in parentheses below indicate the number of independent experiments with at least 40 animals each.

**B)** Estimated breakpoints of NIL59, the smallest interval containing QTL1. Top panel shows gene models near the right breakpoint of NIL59, indicated with blue bar. The right breakpoint lies between a polymorphism in *gcn-2* (indicated with blue asterisk), and a polymorphism in the 3’UTR of *rict-1*, and thus does not include the *rict-1* coding region. Bottom panel shows all CB4856 polymorphisms in *rict-1*, with missense mutations indicated in black text.

**C)** Dauers formed at 27°C by N2, the WN225 NIL, and strains in which the indicated polymorphisms in CB4856 *rict-1* coding sequences have been introduced into N2 *rict-1* sequences via gene editing. Each dot indicates the average number of dauers formed in a single assay. Horizontal bar indicates median. Light-grey thick and thin vertical bars indicate Bayesian 95% and 75% credible intervals, respectively. Numbers in parentheses below indicate the number of independent experiments with at least 40 animals each. *P*-values are relative to N2 (ANOVA with Dunnett-type multivariate-t post-hoc adjustment).

## SUPPLEMENTAL TABLE CAPTIONS

**Table S1.** Strains used in this study.

**Table S2.** Type-I and type-II error rates indicate Bayesian GLMM outperforms frequentist methods for variable dauer data.

